# A partial C_4_ photosynthetic biochemical pathway in rice

**DOI:** 10.1101/2020.06.03.133058

**Authors:** Hsiang-Chun Lin, Stéphanie Arrivault, Robert A Coe, Shanta Karki, Sarah Covshoff, Efren Bagunu, John E. Lunn, Mark Stitt, Robert T. Furbank, Julian M. Hibberd, W Paul Quick

## Abstract

Introduction of a C_4_ photosynthetic pathway into C_3_ rice (*Oryza sativa*) requires installation of a biochemical pump that concentrates CO_2_ at the site of carboxylation in modified bundle sheath cells. To investigate the feasibility of this, we generated a quadruple line that simultaneously expresses four of the core C_4_ photosynthetic enzymes from the NADP-malic enzyme subtype, phospho*enol*pyruvate carboxylase (*Zm*PEPC), NADP-malate dehydrogenase (*Zm*NADP-MDH), NADP-malic enzyme (*Zm*NADP-ME) and pyruvate phosphate dikinase (*Zm*PPDK), in a cell-specific manner. This led to enhanced enzyme activity but was largely neutral in its effects on photosynthetic rate and growth. Measurements of the flux of ^13^CO_2_ through photosynthetic metabolism revealed a significant increase in the incorporation of ^13^C into malate, consistent with increased fixation of ^13^CO_2_ via PEP carboxylase in lines expressing the maize PEPC enzyme. We also showed ^13^C labelling of aspartate indicating additional ^13^CO_2_ fixation into oxaloacetate by PEPC and conversion to aspartate by the endogenous aspartate aminotransferase activity. However, there were no significant differences in labelling of 3-phosphoglycerate (3PGA) or phospho*enol*pyruvate (PEP) indicating limited carbon flux through C_4_ enzymes into the Calvin-Benson cycle. Crossing the quadruple line with a line with reduced glycine decarboxylase H-protein (*Os*GDCH) abundance led to a photosynthetic phenotype characteristic of the reduced *Os*GDCH line and higher labelling of malate, aspartate and citrate. While Kranz anatomy or other anatomical modifications have not yet been installed in these plants to enable a fully functional C_4_ cycle, these results demonstrate for the first-time flux through the carboxylation phase of C_4_ metabolism in transgenic rice containing the key metabolic steps in the C_4_ pathway.

## Introduction

A major recent research objective has been the engineering of a C_4_ photosynthetic pathway into rice (https://C4rice.com; Kajala *et al*., 2011; von Caemmerer *et al*. 2012; Ermakova *et al*. 2019), potentially leading to an increase in radiation use efficiency and yield of up to 50% (Hibberd *et al*. 2008). The C_4_ pathway represents a complex combination of both biochemical and anatomical adaptations that suppresses photorespiration by effectively saturating ribulose bisphosphate carboxylase/oxygenase (Rubisco) with CO_2_. In the majority of C_4_ plants, this is achieved by compartmentalization of photosynthetic reactions between two morphologically distinct cell types: the mesophyll cells (MCs) and the bundle sheath cells (BSCs). Operating across these cells is a biochemical CO_2_ pump elevating the CO_2_ concentration in the BSCs where Rubisco is located (Hatch 1987).

There are three primary variants of this pump characterized by the main decarboxylase reaction (Hatch *et al*. 1975). The NADP-ME subtype was chosen for engineering C_4_ photosynthesis into rice as it is well-characterized in the C_4_ model crop species maize (*Zea mays*) and potentially requires the fewest biochemical enzymes among all C_4_ subtypes (Kajala *et al*., 2011; Weber and von Caemmerer, 2010; Ermakova *et al*. 2019). Each molecule of CO_2_ entering the cytosol of the MCs is first converted to bicarbonate (HCO_3_^-^) by the activity of carbonic anhydrase (CA) and then incorporated into phospho*enol*pyruvate (PEP) by PEP carboxylase (PEPC), yielding the C_4_ acid oxaloacetate (OAA). OAA is taken up into the chloroplast of the MCs where it is reduced to malate by the NADP-dependent malate dehydrogenase (NADP-MDH). Malate is exported back to the cytosol and then diffuses into BSCs through plasmodesmata along a steep concentration gradient. In the BSCs, malate is transported into the chloroplast by an unknown transporter and oxidatively decarboxylated by NADP-dependent malic enzyme (NADP-ME), yielding CO_2_, NADPH and pyruvate. CO_2_ is assimilated by Rubisco, yielding two molecules of 3PGA, about half of which is reduced to triose-phosphate (TPs) using the NADPH provided by NADP-ME in the BSC chloroplast to regenerate RuBP in the Calvin-Benson cycle. The other half of the 3PGA moves to the MCs for reduction to TP in the MC chloroplast and then returns to the BSCs to enter the Calvin-Benson cycle. Pyruvate moves from the BSCs into the chloroplasts of the MCs where it is converted to PEP by pyruvate:phosphate dikinase (PPDK).

In this study, we report on the introduction of part of this biochemical pathway into rice. It has previously been shown that genomic sequences encoding C_4_ proteins give stronger expression in rice than cDNAs (Matsuoka *et al*. 1994). Therefore, we decided to express individual full-length genes of *Zm*PEPC, *Zm*PPDK, *Zm*NADP-MDH and *Zm*NADP-ME (including promoters, untranslated regions, exons and introns) from maize (Kajala *et al*. 2011) in a bid to achieve a C_4_-like pattern of C_4_ gene expression, enzyme localization, enzyme activity and enzyme kinetic properties (Miyao 2003, 2011). Individual lines were then crossed to generate a plant overexpressing all four of these core C_4_ cycle enzymes to investigate the feasibility of installing a functional C_4_ biochemical pathway into rice. This quadruple transgenic line was also crossed with a line with decreased *Os*GDCH protein (Lin *et al*. 2016), and this quintuple line was used to test whether lowering GDC in the MCs of rice primes the plant for introduction of a C_4_ cycle (Sage 2004; Sage *et al*. 2012). We investigated the effect on plant growth and photosynthesis.

To evaluate photosynthetic functionality, we used ^13^CO_2_ labelling experiments (Arrivault *et al*. 2017), similar in concept to the radiolabelling experiments originally performed to characterize flux in C_4_ photosynthesis (Hatch *et al*. 1967, Hatch 1971). Flux of ^13^CO_2_ through photosynthetic metabolism, in particular into C_4_ acids, was determined for the quadruple and quintuple lines, compared to untransformed controls. We show that there was increased labelling of C_4_ acids in both sets of plants compared to wild type, consistent with partial low-level function of a portion of the C_4_ pathway.

## Materials and Methods

### Plant materials

Individual transgenic lines were generated overexpressing four of the core C_4_ cycle enzymes required for a functional NADP-ME C_4_ cycle (Supplementary Figure 1), *Zm*PEPC, *Zm*NADP-MDH, *Zm*NADP-ME and *Zm*PPDK. To express high levels of *Zm*PEPC (GRMZM2G083841), *Zm*PPDK (GRMZM2G306345), *Zm*NADP*-*MDH (GRMZM2G129513) and *Zm*NADP-ME (GRMZM2G085019) in rice, full-length genomic fragments encompassing the genes encoding these maize enzymes and their promoters were cloned into the pSC0 vector (GenBank, Accession no. KT365905; Lin *et al*. 2016). Generation of pSC0/*Zm*PEPC vector was previously described (Giuliani *et al*. 2019a). A pSC0/*Zm*PPDK vector containing a full-length genomic fragment was created by subcloning *Zm*PPDK from pIG121Hm/PPDK (a gift from Mitsue Miyao, NIAS, Japan; Matsuoka 1990) into pSC0. Gibson assembly (Giboson *et al*. 2009) was used to insert the gene into the pSC0 vector. The necessary amplicons from the pIG121Hm/PPDK and pSC0 templates were amplified using Primer I: 5’-ATGCTCAACACATGAGCGAAGGGCCCATGACCATGATTACGCCAAG, Primer II: 5’-TGTGCATGTCGCTAGGATCCGGTACCGAATGCTAGAGCAGCTTGA, Primer III: TCAAGCTGCTCTAGCATTCGGTACCGGATCCTAGCGACATGCACA, Primer IV: 5’-CTTGGCGTAATCATGGTCATGGGCCCTTCGCTCATGTGTTGAGCAT). The full-length genomic sequences of *Zm*NADP*-*ME and *Zm*NADP*-*MDH were amplified from bacterial artificial chromosomes sourced from BACPAC resources (Children’s Hospital Oakland, California) (Coordinates CH201-14H23 for *Zm*NADP-ME and CH201-117G14 for *Zm*NADP*-*MDH) by PCR using primers (5’-ACGACGGCCAGTGCCAAGCTTCCCTTCCGTCAGCAGATTAGGCG and 5’-ATTATTATGGAGAAACTCGAGGCAACATGGTTCTGGACCGATTCAG for *Zm*NADP*-* MDH;5’-ACGACGGCCAGTGCCAAGCTTGGAATGACCACGAAATCGTCAAGCTAATCC and 5’-ATTATTATGGAGAAACTCGAGCTGTTACTGCTCTTTCCACTACTGAAGCAG for *Zm*NADP*-*ME) and subcloned into pSC0 vector. A binary vector with the hygromycin B resistance gene, pCAMBIA1300, was co-transformed with these vectors to allow for selection. The transformation of *indica* rice (*Oryza sativa* cv. IR64) was performed at the International Rice Research Institute (IRRI; Los Baños, Philippines) following a previously described method (Lin *et al*. 2016). In almost all cases, three independent single insertion homozygous transgenic lines with high transgene expression were selected for molecular and biochemical evaluation. However, for *Zm*NADP-ME, protein expression was only detected in a single transgenic line containing >6 copies of the overexpression construct and so this was the only line that could be taken forward. The line with highest protein expression for each gene was selected for stacking into a multi-transgenic single line through conventional crossing to create a quadruple cross line (PEPC/PPDK/MDH/ME) for investigation in the present study. The *Zm*PEPC and *Zm*NADP*-*MDH single transgenic lines were initially crossed to create a double transgene line that was then crossed with the *Zm*PPDK single transgenic line. This triple line (PEPC/PPDK/MDH) was then crossed with the *Zm*NADP*-*ME single transgenic line to produce a quadruple line (PEPC/PPDK/MDH/ME). A quintuple cross (PEPC/PPDK/MDH/ME/*gdch*) was then generated by crossing the quadruple F_2_ line with a single *Osgdch* knockdown line (*gdch*-38) described by Lin *et al*. (2016). The presence of transgenes was determined by genomic PCR and protein accumulation by immunoblotting in each crossed line.

### Plant growth

Plants were grown under natural light conditions in a screenhouse with a day/night temperature of 35/28 ± 3°C at the International Rice Research Institute (Los Baños, Philippines: 14° 10019.900N, 121° 15022.300E). Maximum irradiance was 2000 µmol photons m^-2^ s^-1^ on a sunny day. Plants were grown in 7-liter pots filled with soil from the IRRI upland farm.

### Immunoblotting

Leaf samples for soluble protein extraction were harvested from the fourth fully-expanded leaf at the mid-tillering stage between 09:00 h and 11:00 h, and stored on ice immediately. Leaves were homogenized to a fine powder using a nitrogen-cooled mortar and pestle. Proteins were extracted and fractionated by SDS-PAGE as described previously (Lin *et al*. 2016). Samples were loaded based on equal leaf area (0.2364 mm^2^ for *Zm*PEPC and *Zm*PPDK, and 2.364 mm^2^ for *Zm*NADP*-*MDH, *Zm*NADP*-* ME and *Os*GDCH). After electrophoresis, proteins were electroblotted onto a polyvinylidene difluoride membrane and probed with rabbit antisera against *Zm*PEPC, *Zm*NADP*-*MDH, *Zm*NADP*-*ME (all provided by Richard Leegood, University of Sheffield, UK), *Zm*PPDK (provided by Chris Chastain, Minnesota State University, USA) and *Os*GDCH protein (provided by Asaph Cousins, Washington State University, USA). The dilutions of *Zm*PEPC, *Zm*PPDK, *Zm*NADP*-*MDH, *Zm*NADP*-*ME, and *Os*GDCH antisera were 1:20,000, 1:20,000, 1:5,000, 1:2,000 and 1:100, respectively. A peroxidase-conjugated goat anti-rabbit IgG secondary antibody (Sigma-Aldrich, USA; https://www.sigmaaldrich.com/) was used at a dilution of 1:5,000 and immunoreactive bands were visualized with ECL Western Blotting Detection Reagents (GE Healthcare, UK; https://www.gelifesciences.com).

### Immunolocalization

The middle portion of the seventh fully expanded leaf at the mid-tillering stage was sampled between 09:00 h and 11:00 h and processed as described previously by Lin *et al*. (2016). After fixation and cutting, the thin leaf sections were probed with the antisera against *Zm*NADP*-*MDH, *Zm*NADP*-*ME, *Zm*PPDK and *Zm*PEPC at dilutions of 1:500, 1:25, 1:10 and 1:200, respectively. The secondary Alexa Fluor 488 goat anti-rabbit IgG (Invitrogen, USA; https://www.thermofisher.com/ph/en/home/brands/invitrogen.html) antibody was used at a dilution of 1:200. The sections were visualized on a BX61 microscope fitted with a Disk Scanning Unit attachment microscope (Olympus, USA; https://www.olympus-global.com/) with fluorescence function under DAPI, RFP and GFP filters.

### Enzyme activity measurement

Leaf samples were harvested between 09:00 h and 11:00 h from the youngest fully-expanded leaf of plants at the mid-tillering stage, and frozen immediately. Leaves were homogenized to a fine powder using a nitrogen-cooled mortar and pestle and extracted in 250 μL of buffer containing: 50 mM HEPES-KOH, pH7.4, 5 mM MgCl_2_, 1 mM EDTA, 1 mM Dithiothreitol, 1% (v/v) glycerol. After centrifugation at 10,000x*g* for 2 min at 4°C, the supernatant was collected for enzyme activity measurements. PEPC enzyme activity was assayed using a method modified from Meyer *et al*. (1988) and Ueno *et al*. (1997). The PEPC reaction mixture contained: 100 mM HEPES-NaOH, pH 7.5, 10 mM MgCl_2_, 1 mM NaHCO_3_, 5 mM G6P, 0.2 mM NADH, 12 unit/mL MDH (from pig heart; Roche Diagnostics, Basel, Switzerland; https://www.roche.com/), and the reaction was started by adding PEP to a final concentration of 4 mM. PPDK enzyme activity was assayed as described by Fukayama *et al*. (2001). NADP-MDH activity was determined by a method modified from Tscuhida *et al*. (2001). NADP-MDH reaction mixture contained: 50 mM HEPES-KOH, pH 8, 70 mM KCl, 1 mM EDTA, 1 mM DTT, and 0.2 mM NADPH, and the reaction was started by adding OAA to a final concentration of 1 mM. NADP-ME activity was measured by a method modified from Tscuhida *et al*. (2001; protocol 1). The activities of PEPC, PPDK, NADP-MDH and NADP-ME were measured spectrophotometrically at 340 nm at 25°C, 30°C, 25°C and 25° C, respectively.

### Gas exchange measurements

Leaf gas-exchange measurements were made using a Li-6400XT infrared gas analyzer (LI-COR Biosciences, USA; https://www.licor.com/) fitted with a standard 2 × 3 cm leaf chamber and a 6400-02B light source. Measurements were made at a constant airflow rate of 400 μmol s^-1^, leaf temperature of 25°C, leaf-to-air vapor pressure deficit between 1.0 and 1.5 kPa and relative humidity of 60-65%. Data were acquired between 08:00 h and 13:00 h. Measurements were made from two youngest fully expanded leaves for each plant during the tillering stage. The mid-portions of leaves were acclimated in the cuvette for approximately 30 min before measurements were made. The response curves of the net CO_2_ assimilation rate (*A*, µmol m^-2^ s^-1^) to changing intercellular *p*CO_2_ concentration (*Ci*, µmol CO_2_ mol air^-1^) were acquired by decreasing *Ca* (*p*CO_2_ concentration in the cuvette) from 2000 down to 20 µmol CO_2_ mol air^-1^ at a photosynthetic photon flux density (PPFD) of 1500 or 2000 µmol photon m^-2^ s^-1^. The CO_2_ compensation point (Γ) and maximum carboxylation efficiency (*CE*) were calculated from the intercept (Vogan *et al*. 2007) and slope (Wang *et al*. 2006) of the CO_2_ response curves. Light response curves were acquired by increasing the PPFD from 0 to 2000 µmol photon m^-2^ s^-1^ at *Ca* 400 µbar. The quantum efficiency for CO_2_ assimilation (Φ) and respiration rates (*R*_*d*_) were calculated from the slope and intercept of the light-response curves (PPFD < 100 μmol photons m^-2^ s^-1^).

### ^13^CO_2_ pulse-labelling and quenching procedure

Carbon flux analysis was performed with a custom-built gas exchange freeze clamp apparatus (Supplementary Figure 2). Measurements were made from two youngest fully expanded leaves for each plant during the tillering stage. Two leaves of up to 22 cm in length were placed inside a gas exchange chamber (23.5 × 4.5 × 0.4 cm), the top was constructed from a piece of clear flexible plastic to allow light penetration and the bottom from a sheet of aluminum foil to accelerate cooling when freeze clamping. The foil and plastic were attached with foil tape to a three-sided aluminum frame to provide rigidity. Two holes were drilled through the side of the frame to accommodate the air inlet and outlet tubes. A third hole on the end enabled thermocouples to be threaded through the frame to measure leaf and air temperature inside the labelling chamber. The chamber was then placed in a mounting frame allowing the leaf to be inserted prior to sealing the chamber with a foam gasket secured with bulldog clips. The mounting frame was positioned horizontally between two LED banks capable of providing illumination to the upper leaf surface of up to 1,000 µmol photons m^2^ s^-1^.

Air was drawn from outside the laboratory through a compressor. The air stream passed through an oil water separator and flow control valve into a copper coil placed in an ice bath for cooling. The air stream could be directed into the leaf chamber or by-passed into a CO_2_ conditioning unit. In the latter, CO_2_ could be removed from the air with soda lime and then optionally enriched with ^13^CO_2_ gas (300 ppm) prepared by mixing NaH^13^CO_3_ (Sigma-Aldrich, USA; https://www.sigmaaldrich.com/) with 2-hydroxypropanoic acid (lactic acid). The flow of air passing over the leaf (3 ml/min) was adjusted with a flow controller, and the CO_2_ concentration of the incoming and outgoing air streams measured with two CO_2_ analyzers (WMA-5, PP-Systems, USA; https://ppsystems.com/). The humidity of the air inside the chamber was maintained at ∼60% with the addition of water to the soda lime chamber or reduced by passing air through a chamber of silica gel.

The leaf chamber was mounted on the stand in such a way that the plane of the leaf was halfway between two pneumatically operated aluminum bars. These were cooled with liquid nitrogen, and when released they clamped together fitting inside the aluminum chamber frame, very rapidly freezing the leaf. A fan was mounted horizontally to the bars to blow the fog from the liquid nitrogen away from the chamber to ensure there was minimal disruption to the environment before freezing occurred. The lower bar was positioned in such a way as to push the chamber up on closure. The cold temperatures and force of the bars closing meant the chamber disintegrated enabling the leaf to be removed with tweezers and placed in a liquid nitrogen bath for 10 s before subsequent storage at −80 °C. To perform ^13^CO_2_ pulse-labelling, leaves were acclimated at steady-state conditions prior to scrubbing CO_2_ from the incoming air stream, then subjected to ^13^CO_2_ enriched air for a duration of 0 and 60 s and metabolic activity was quenched at these time points by freeze clamping the leaves as above. Freeze-quenched tissue was ground into a fine powder by mortar and pestle in liquid nitrogen. Finely-ground leaf tissues were freeze-dried for 3 days and placed into sealed tubes. The sealed tubes containing finely-ground lyophilized leaf tissue samples were shipped to Max Planck Institute of Molecular Plant Physiology, Germany for the metabolite analyses.

### Metabolite analyses and calculation of total pool size, enrichment and isotopomer distribution

Aliquots (3 or 5 mg) of finely-ground lyophilized rice leaf tissue were extracted with chloroform-methanol as described in Arrivault *et al*., (2017), and the lyophilized extracts were resuspended in 300 µL or 600 µL purified (Millipore, USA) water, respectively. Isotopomers were measured by reverse-phase LC-MS/MS (malate, aspartate, 3PGA, PEP, citrate+isocitrate; Arrivault *et al*. 2017; n.b. citrate & isocitrate were not resolved using this method) and anion-exchange LC-MS/MS (malate, PEP, citrate; Lunn *et al*. 2006 with modifications as described in Figueroa *et al*. 2016). Total amounts of malate, aspartate, citrate+isocitrate and citrate were calculated by summing isotopomers. The total amounts of 3PGA and PEP were determined enzymatically in trichloroacetic acid extracts using a Sigma-22 dual-wavelength photometer (Merlo *et al*. 1993), with PEP being measured in freshly prepared extracts. ^13^C enrichment and relative isotopomer distribution were calculated as in Szecowka *et al*. (2013).

### Statistical analysis

Statistical analysis for all experiments was performed in R version 3.0.0 (The R Foundation for Statistical Computing, Vienna, Austria) using a one-way analysis of variance (ANOVA) and a Tukey post-hoc test or a Student’s t-test with a *p*-value of <0.05.

## Results

### Overexpression of C_4_ cycle genes in *Oryza sativa*

Immunoblotting of F_2_ generation transgenic plants showed that the protein of the correct size for *Zm*PEPC, *Zm*NADP-MDH, *Zm*NADP-ME and *Zm*PPDK was stably expressed in both the quadruple and quintuple cross rice lines, with *Os*GDCH protein almost undetectable in the quintuple cross (Figure 1). In all lines, protein abundance was lower than that of wild-type maize plants. We next sought to determine whether these proteins were localized to the correct cell type and subcellular compartment. Immunolocalization analysis of the quadruple cross line revealed that *Zm*PEPC was localized to the cytosol of MCs similar to the single *Zm*PEPC transgenic line (Giuliani *et al*. 2019b) and *Zm*PPDK, was localized to the chloroplast in both BSCs and MCs (Supplementary Figure 3). The *Zm*NADP-MDH and *Zm*NADP-ME antisera cross-reacted with native protein in wild-type rice and so it was not possible to distinguish protein encoded by the endogenous rice gene from that encoded by the maize transgene. Overexpression of these enzymes conferred enhanced enzyme activity (Supplementary Table 1). In the quadruple cross line, PEPC activity was 18.4-fold higher compared to wild-type rice plants, PPDK 5.8-fold higher, NADP-MDH 9.9-fold and NADP-ME 4.1-fold.

### Phenotypic and photosynthetic perturbations associated with overexpression of C_4_ cycle genes

Given that protein levels and activities of all four introduced C_4_ enzymes were enhanced compared to wild-type plants, we investigated whether this affected growth and photosynthesis. None of the crossed lines consistently showed altered chlorophyll content (Figure 2A). Tiller number in the quintuple cross lines (Figure 2B) and plant height in both quadruple and quintuple crosses (Figure 2C) were significantly reduced. Phenotypic perturbations were most marked in the quintuple cross lines (Supplementary Figure 4), although the plants still developed and flowered at the same time with wild-type plants. These growth perturbations were not observed in the single C_4_ gene transgenic lines of *Zm*PEPC, *Zm*PPDK, *Zm*NADP*-*MDH and *Zm*NADP*-*ME (Giuliani et al. 2019) but were observed in the single *Osgdch* knockdown line (Lin *et al*. 2016).

To investigate whether overexpression of C_4_ genes impacted photosynthesis, the response of net CO_2_ assimilation rate (*A*) to CO_2_ concentration under photorespiratory conditions (21% O_2_) was measured. In the quadruple cross line there were no differences in CO_2_ assimilation (Figure 3A, C), Γ, CE, *R*_*d*_ or Φ compared to wild-type plants (Table 1). These results suggested that enhanced activity of C_4_ cycle enzymes in a cell-specific manner does not significantly affect leaf level photosynthetic gas-exchange and that the phenotypic perturbations are not associated with reduced CO_2_ assimilation.

For the quintuple cross, the CO_2_ assimilation rate in response to increasing intercellular CO_2_ concentrations (*Ci*) was reduced, most notably at lower CO_2_ concentrations (<700 µmol CO_2_ mol^-1^, Figure 3B). Under non-photorespiratory conditions (2% O_2_, Figure 3B) CO_2_ assimilation rates were similar to wild-type plants. Consistent with this, the quintuple cross had a significantly higher Γ under high photorespiratory conditions but not under low photorespiratory conditions (Table 1). In response to changes in photon flux density, photosynthesis was saturated at lower light levels than in wild-type plants (400 µmol photons m^-2^ s^-1^ versus 1750 µmol photons m^-2^ s^-1^, respectively, Figure 3D) with significantly lower Φ and higher *R*_*d*_ than wild-type plants (Table 1).

To investigate these photosynthetic responses for the quintuple cross line in more detail, CO_2_ responses were measured under conditions conducive to low and high rates of photorespiration. Under low light (400 µmol photons m^-2^ s^-1^) and high CO_2_ (2000 µmol CO_2_ mol air^-1^), conditions conducive to low photorespiration, CO_2_ responses of the quintuple cross line were similar to wild-type plants (Supplementary Figure 5A, D). Under high light (1500 µmol photons m^-2^ s^-1^) and low CO_2_ (400 µmol CO_2_ mol air^-1^), conditions conducive to high photorespiration, CO_2_ assimilation was lower in the quintuple cross (Supplementary Figure 5B, C). Correspondingly, Γ was higher, and carboxylation efficiency (*CE*) lower under conditions conductive to high rates of photorespiration but not under non-photorespiratory conditions (Supplementary Table 2). This is consistent with the photorespiratory-deficient phenotype observed in the single *Osgdch* knockdown lines (Lin *et al*. 2016).

### Increased incorporation of ^13^C into C_4_ acids associated with overexpression of C_4_ cycle enzymes

We performed experiments to measure the flux of ^13^CO_2_ through photosynthetic metabolism to investigate whether there was partial functionality of a C_4_ pathway in these plants. There was significantly more incorporation of ^13^C into malate in the quadruple and quintuple cross lines than in wild-type plants (Figure 4, Supplementary Figure 6), with the *m*_*1*_ isotopomer being more abundant than the other ^13^C-labelled isotopomers (Table 2), consistent with increased fixation of ^13^CO_2_ via PEPC in the transgenic lines expressing the maize PEPC enzyme. Labelling of aspartate was also significantly higher than wild-type in the quintuple line (Figure 4, Supplementary Figure 6), with the *m*_*1*_ isotopomer being more abundant that *m*_*2*_*-m*_*4*_ isotopomers (Table 2), indicating additional ^13^CO_2_ fixation into oxaloacetate by maize PEPC and conversion to aspartate by endogenous rice aspartate aminotransferase activity. There was almost complete labelling of the 3PGA pool in wild-type plants after a 60 s pulse, and similarly high levels of labelling of 3PGA were observed in the quadruple and quintuple lines (Figure 4, Supplementary Figure 6). There was also substantial labelling of PEP after a 60 s pulse, with no significant differences between the transgenic lines and wild-type plants (Figure 4, Supplementary Figure 6). There was almost no labelling of citrate and isocitrate in wild-type plants and the quadruple line (Figure 4). In contrast, the enrichment of ^13^C in citrate and isocitrate in the quintuple line was 10-fold higher than in wild-type plants.

## Discussion

We have previously shown that overproduction of individual C_4_ enzymes in rice has no consistent effect on CO_2_ assimilation or plant growth (Giuliani *et al*. 2019b). The exception to this was the transgenic line overexpressing *Zm*NADP-ME which exhibited a small decrease in plant height and reduced maximal photosynthetic rate at high CO_2_. Previous attempts to overproduce *Zm*NADP-ME in rice have led to increased photoinhibition of photosynthesis, leaf chlorophyll bleaching and serious stunting attributed to an increase in the NADPH/NADP^+^ ratio in the chloroplast stroma due to the exchange with 2-oxoglutarate involved in photorespiration (Tsuchida *et al*. 2001). Severe phenotypic effects were not observed in our *Zm*NADP-ME line; however, we were only able to advance a single line containing 6 copies of the construct in which protein accumulation was higher than in our rice control, but still only around 10% of the activity found in maize. In contrast, the experiments of Tsuchida *et al*. (2001) used a rice chlorophyll a/b binging protein (*cab*) promoter allowed for high level but not cell specific expression, and activities of up to 60% of maize levels were achieved, leading to a much more severe phenotype. Overproduction of all four targeted C_4_ enzymes in a single plant led to a slight decrease in tiller number and plant height but otherwise growth and photosynthesis were unaffected. These results are consistent with previous reports of engineering a single-cell C_4_ pathway in rice (Taniguchi *et al*. 2008). A quintuple cross that combined over-expression of the four C_4_ enzymes, *Zm*PEPC, *Zm*NADP-MDH, *Zm*NADP-ME and *Zm*PPDK with knockdown of the native rice *Os*GDCH, thereby compromising the photorespiratory pathway, led to further reductions in tiller number and plant height. A strong negative effect on photosynthesis was also observed in the quintuple cross consistent with the photorespiratory-deficient phenotype of the single *Osgdch* knockdown line (Lin *et al*. 2016).

Our results show that *Zm*PEPC is catalytically active *in vivo* when expressed in combination with other C_4_ enzymes in rice, and substantially increases the fixation of CO_2_ into C_4_ acids. This is in contrast to published radiolabelling studies of rice expressing *Zm*PEPC alone (Fukayama *et al*. 2003; Miyao *et al*. 2011) in which there was no increase in incorporation of labelled carbon into C_4_ acids, despite the extractable activity of PEPC in these plants approaching or exceeding maize levels. Despite strong evidence for operation of the C_4_ pathway in our transgenic lines up to the point of malate production, there was no evidence for the regeneration of PEP via the rest of the C_4_ cycle. The labelling of PEP at a similar level in all three genotypes, wild-type, quadruple and quintuple crosses, suggests that rather than being produced by a functional C_4_ cycle, PEP is being produced from 3PGA via 2PGA catalyzed by phosphoglyceromutase and enolase (Furbank and Leegood 1984), consistent with the majority of CO_2_ still being fixed via Rubisco in C_3_ photosynthesis rather than through the operation of a complete C_4_ cycle.

The very low incorporation of ^13^C into citrate and isocitrate in wild-type rice plants is consistent with previous studies (Tcherkez *et al*. 2009; Szecowka *et al*. 2013) indicating little flux of carbon into the tricarboxylic acid (TCA) cycle via mitochondrial pyruvate dehydrogenase (mPDH) in the light, due to deactivation of the mPDH by phosphorylation (Randall *et al*. 1996; Tovar-Méndez *et al*. 2003). The increased labelling of citrate in the quintuple cross line suggests that the mPDH is more active in the light in this line, potentially leading to respiration of C_4_ acids via the TCA cycle, which would be deleterious for C_4_ photosynthetic flux. This might be due to lower rates of photorespiration leading to less photophosphrylation of PDH (Tovar-Méndez *et al*. 2003). Further, increased levels of pyruvate in the mitochondria (from decarboxylation of malate by NAD-malic enzyme), can inhibit the mPDK kinase (Schuller & Randall 1990). We propose that there is a modified regulation of the TCA cycle to avoid wasteful respiration of C_4_ acids, and that such modification might have been needed for the evolution of an efficient C_4_ photosynthetic pathway.

Evidence that *Zm*PEPC can be localized to the cytosol of MCs (Giuliani *et al*. 2019b) and is catalytically active, leading to the fixation of CO_2_ into C_4_ acids provides important evidence in support of installing a fully functional C_4_ photosynthetic pathway into rice. However, absolute quantification of flux into and through C_4_ acids would require further pulse-chase labelling studies and may prove difficult with the low rates of labelling relative to C_3_ photosynthetic fixation obtained in the current transgenic lines.

Cell specificity of expression of the C_4_ enzymes introduced into the rice lines shown here remains an important issue. We have been able to achieve localization of *Zm*PPDK to rice MCs and additional localization to the chloroplast of rice BSCs by the use of a native *Zm*PPDK promoter. This result is consistent with *Zm*PPDK expression in both MCs and BSCs in maize (Sheen and Bogorad 1987; Miyao et al. 2011). However, we were unable to conclude with confidence whether there was correct localization of *Zm*NADP-ME and *Zm*NADP-MDH in BSCs and MCs, respectively. When used to drive β*-GLUCURONIDASE* (*GUS*) expression, the promoter used for *Zm*NADP-ME expression was shown to lead to GUS accumulation in both BSCs and MCs in rice (Nomura *et al*. 2005), suggesting that exclusive localization of NADP-ME in the BSCs of our plants is unlikely. Incorrect or partial localization of enzymes would potentially limit the operation of a C_4_ cycle (Miyao *et al*. 2011), and higher-level expression of *Zm*NADP-ME than that achieved in our lines, particularly if mis-expressed in the M cells, may produce quite deleterious phenotypes (Tsuchida *et al*. 2001).

In addition to high level, cell specific expression of C_4_ cycle enzymes in rice, fully functional C_4_ photosynthetic biochemistry requires appropriate enzyme regulation in the environment of a rice leaf cell (Burnell and Hatch 1985; Chastain *et al*. 1997). For example, the activity of C_4_ specific PPDK is regulated in the light through protein phosphorylation by the PPDK regulatory protein (Burnell and Hatch 1985). Similarly NADP-MDH is regulated by light through the thioredoxin cascade (Miginiac-Maslow *et al*. 2000). Both enzymes are regulated in the same manner even when expressed within C_3_ leaves (Fukayama *et al*. 2001; Taniguchi *et al*. 2008). A recent study has shown that C_4_ NADP-ME is also regulated in the light by reversible phosphorylation at Ser419 which is involved in the binding of NADP at the active site (Boydiloya et al. 2019). In contrast, PEPC is regulated by both metabolite effectors and reversible phosphorylation, but the mechanisms of regulation in C_3_ and C_4_ leaves are different (Vidal and Chollet 1997). Indeed, Fukayama *et al*. (2003) observed inappropriate phosphorylation of PEPC in their transgenic rice lines and proposed this as a reason for lack of labeling of C_4_ acids in the light. The regulatory mechanisms for other enzymes are less well understood. It is unclear at present whether enzyme levels per se or enzyme regulation in our rice transgenic lines, or both, is limiting C_4_ flux.

In addition to the metabolic enzymes of the C_4_ pathway, there is a need to identify and overproduce the metabolite transporters required to support C_4_ photosynthesis (Weber and von Caemmerer 2010; Ermakova *et al*. 2019). While many of the necessary transporters may be present at low levels in C_3_ chloroplasts (Weber and von Caemmerer 2010), their activities might be insufficient to mediate the greater fluxes across the chloroplast envelope that are required for operation of a C_4_ cycle.

A plethora of other changes are required to support a fully functional C_4_ pathway in rice. This includes, but is not limited to, engineering the correct leaf anatomy (Hattersley and Watson 1975, Dengler *et al*. 1994; Muhaidat *et al*. 2007; Dengler and Taylor 2000) and morphological specializations such as increased vein density (Sedelnikova *et al*. 2018). In addition, thought must be given to the photosynthetic functionalisation of the BSCs of rice, which contain a large central vacuole, with very few mitochondria, peroxisomes or chloroplasts (Sage and Sage 2009). Where chloroplasts do occur, they are smaller than those in MCs. In addition, MCs of rice are highly lobed to assist with photorespiratory CO_2_ scavenging (Sage and Sage 2009); whereas the MCs of C_4_ species are not. Increasing chloroplast number and volume in the BSCs will no doubt be important for achieving C_4_ photosynthesis in rice (Chonan 1970, 1978; Dengler *et al*. 1994; Ueno *et al*. 2006; Wang *et al*. 2017). Insufficient chloroplast volume in the BSCs of rice may have led to limitations in C_4_ acid decarboxylation in the transgenic lines described here. Other modifications such as cross-sectional area of the BSCs, modifying the cell wall properties for diffusion of CO_2_ (von Caemmerer and Furbank 2003) and increasing plasmodesmatal frequency at the BSC / MC interface to support metabolite diffusion may be necessary (Ermakova *et al*. 2019). The genetic regulators of many of these changes are not known, and so future goals include identification and incorporation of necessary genes for anatomical modifications into a version of the current biochemical prototype, with the ultimate goal of engineering an efficient C_4_ pathway in rice.

## Supplementary Materials

Supplementary Dataset A. Isotopomer and metabolite amounts, 13C enrichments and relative isotopomer abundances of malate, aspartate, 3PGA, PEP and citrate+isocitrate in wild-type, quadruple and quintuple lines.

Supplementary Dataset B. Isotopomer and metabolite amounts, 13C enrichments and relative isotopomer abundances of malate, aspartate, 3PGA, PEP and citrate in wild-type and quintuple lines.

## Author Contributions

HCL, SA, RAC, JEL, MS, RTF and WPQ designed the experiments together. HCL provided all the plant materials. HCL and EB performed enzyme activity assay, immunobloting, immunolocalization and gas exchange measurements. WPQ, RTF, MS, JEL and RAC designed the gas exchange freeze clamp apparatus. SA performed metabolite analysis. HCL, SA, RAC and WPQ wrote the manuscript. SC and JMH designed constructs. SK performed plant transformation. HCL and RAC performed the ^*13*^*CO*_*2*_ labelling experiment.

## Acknowledgements

This paper is dedicated to the memory of Dr. John Sheehy, who was the architect and inspiration of the C_4_ Rice Consortium and its attempts to engineer a rice plant with C_4_ or C_4_-like photosynthesis. We wish to thank Florencia Montecillo, Juvy Reyes and Irma Canicosa for their help with plant transformation, husbandry and physiological measurements at IRRI C_4_ Rice Center as well as Regina Feil and Manuela Guenther for metabolite measurements at the MPI-MP.

