## Supplemental for "A partial C_4_ photosynthetic biochemical pathway in rice"

***Title***

***Abstract***

Introduction of a C_4_ photosynthetic pathway into C_3_ rice (*Oryza sativa*) requires installation of a biochemical pump that concentrates CO_2_ at the site of carboxylation in modified bundle sheath cells. To investigate the feasibility of this, we generated a quadruple line that simultaneously expresses four of the core C_4_ photosynthetic enzymes from the NADP-malic enzyme subtype, phospho*enol*pyruvate carboxylase (*Zm*PEPC), NADP-malate dehydrogenase (*Zm*NADP-MDH), NADP-malic enzyme (*Zm*NADP-ME) and pyruvate phosphate dikinase (*Zm*PPDK), in a cell-specific manner. This led to enhanced enzyme activity but was largely neutral in its effects on photosynthetic rate and growth. Measurements of the flux of ^13^CO_2_ through photosynthetic metabolism revealed a significant increase in the incorporation of ^13^C into malate, consistent with increased fixation of ^13^CO_2_ via PEP carboxylase in lines expressing the maize PEPC enzyme. We also showed ^13^C labelling of aspartate indicating additional ^13^CO_2_ fixation into oxaloacetate by PEPC and conversion to aspartate by the endogenous aspartate aminotransferase activity. However, there were no significant differences in labelling of 3-phosphoglycerate (3PGA) or phospho*enol*pyruvate (PEP) indicating limited carbon flux through C_4_ enzymes into the Calvin-Benson cycle. Crossing the quadruple line with a line with reduced glycine decarboxylase H-protein (*Os*GDCH) abundance led to a photosynthetic phenotype characteristic of the reduced *Os*GDCH line and higher labelling of malate, aspartate and citrate. While Kranz anatomy or other anatomical modifications have not yet been installed in these plants to enable a fully functional C_4_ cycle, these results demonstrate for the first-time flux through the carboxylation phase of C_4_ metabolism in transgenic rice containing the key metabolic steps in the C_4_ pathway.

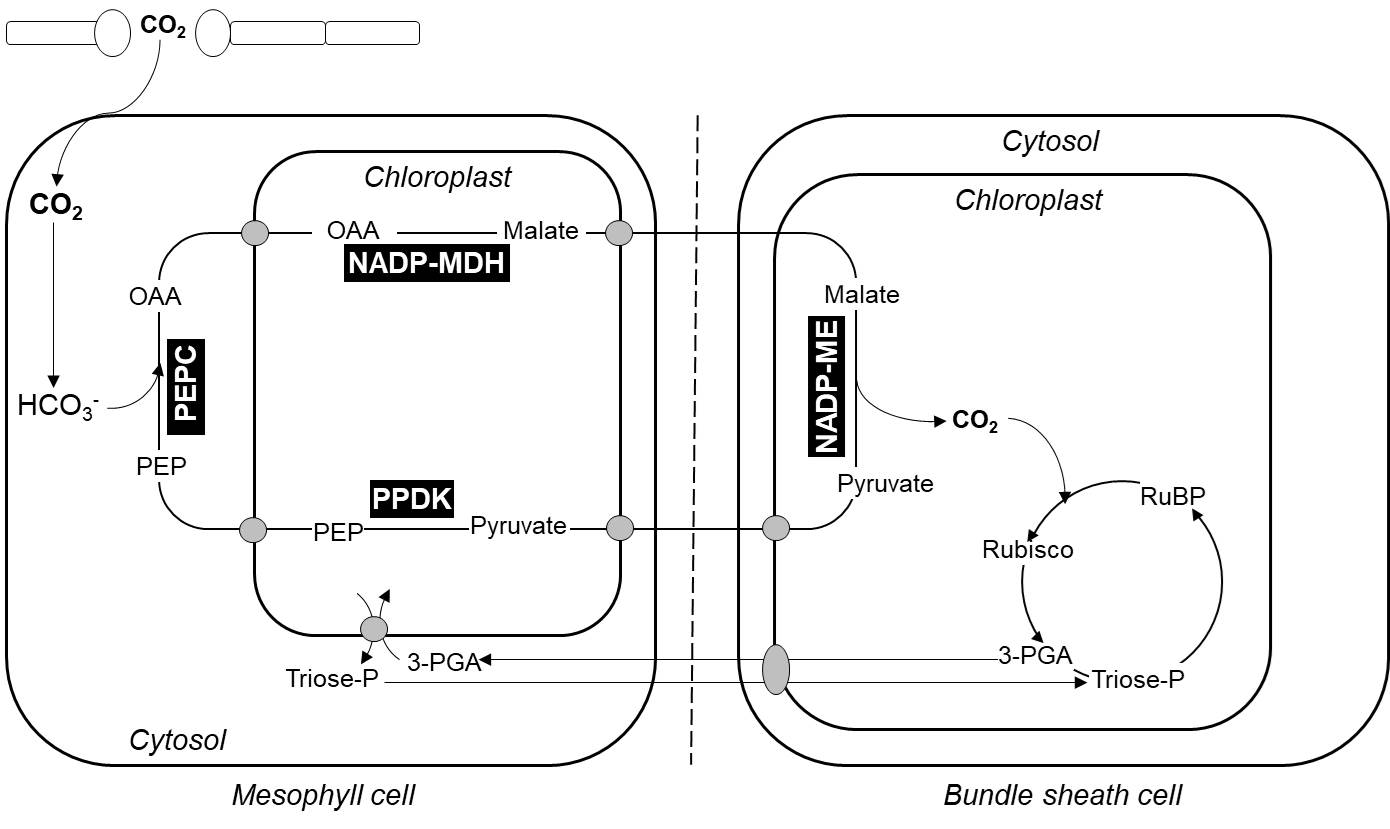

**Supplementary Figure 1.** Schematic representation of the NADP-ME subtype of C_4_ photosynthesis. Carbon dioxide (CO_2_) is converted to bicarbonate (HCO_3_^-^); this is then fixed by phospho*enol*pyruvate carboxylase (PEPC) catalyzing the formation of oxaloacetate (OAA). OAA is reduced to malate by NADP-dependent malate dehydrogenase (NADP-MDH). This is oxidatively decarboxylated by NADP-dependent malic enzyme (NADP-ME), yielding CO_2_, NADPH and pyruvate. CO_2_ is assimilated by Rubisco. Pyruvate is converted to PEP by pyruvate:phosphate dikinase (PPDK).

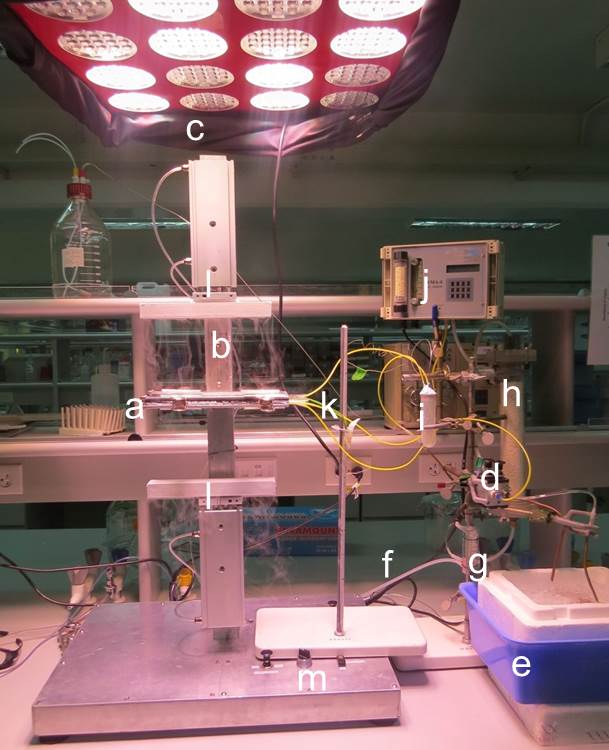

**Supplementary Figure 2.** Custom gas exchange freeze clamp apparatus. (a) custom gas exchange chamber, (b) mounting frame, (c) LED light bank, (d) air conditioning system control, (e) copper coil and ice bath for air cooling, (f) air inlet, (g) flow control, (h) CO_2_ scrubber, (i) ^13^CO_2_ gas delivery system, (j) CO_2_ analyser (second unit not shown), (k) chamber air inlet and outlet (thermocouple not shown), (l) liquid nitrogen cooled aluminium bars, (m) pneumatic control.

**Supplementary Table 1.** The enzyme activities of PEPC, NADP- MDH, NADP-ME and PPDK.

|  | PEPC | | |  | NADP-MDH | | |  | NADP-ME | | |  | PPDK | | |  |
| --- | --- | --- | --- | --- | --- | --- | --- | --- | --- | --- | --- | --- | --- | --- | --- | --- |
|  | µmol s^-1^ m^-2^ | | |  | µmol s^-1^ m^-2^ | | |  | µmol s^-1^ m^-2^ | | |  | µmol s^-1^ m^-2^ | | |  |
| Maize | 42.54 | ± | 1.09 | ^a^ | 12.81 | ± | 0.39 | ^b^ | 24.21 | ± | 2.38 | ^a^ | 9.44 | ± | 1.34 | ^a^ |
| WT | 1.03 | ± | 0.01 | ^c^ | 3.50 | ± | 0.03 | ^c^ | 0.56 | ± | 0.07 | ^c^ | 0.30 | ± | 0.03 | ^c^ |
| Quadruple | 18.96 | ± | 2.56 | ^b^ | 34.98 | ± | 3.26 | ^a^ | 2.34 | ± | 0.10 | ^b^ | 1.76 | ± | 0.14 | ^b^ |

Values are average ± SE of 2-3 plants of maize, wild-type rice (WT) and F_2_ quadruple crosses. Different letters within groups indicated those values that are statistically different based on a one-way ANOVA, P-value<0.05.

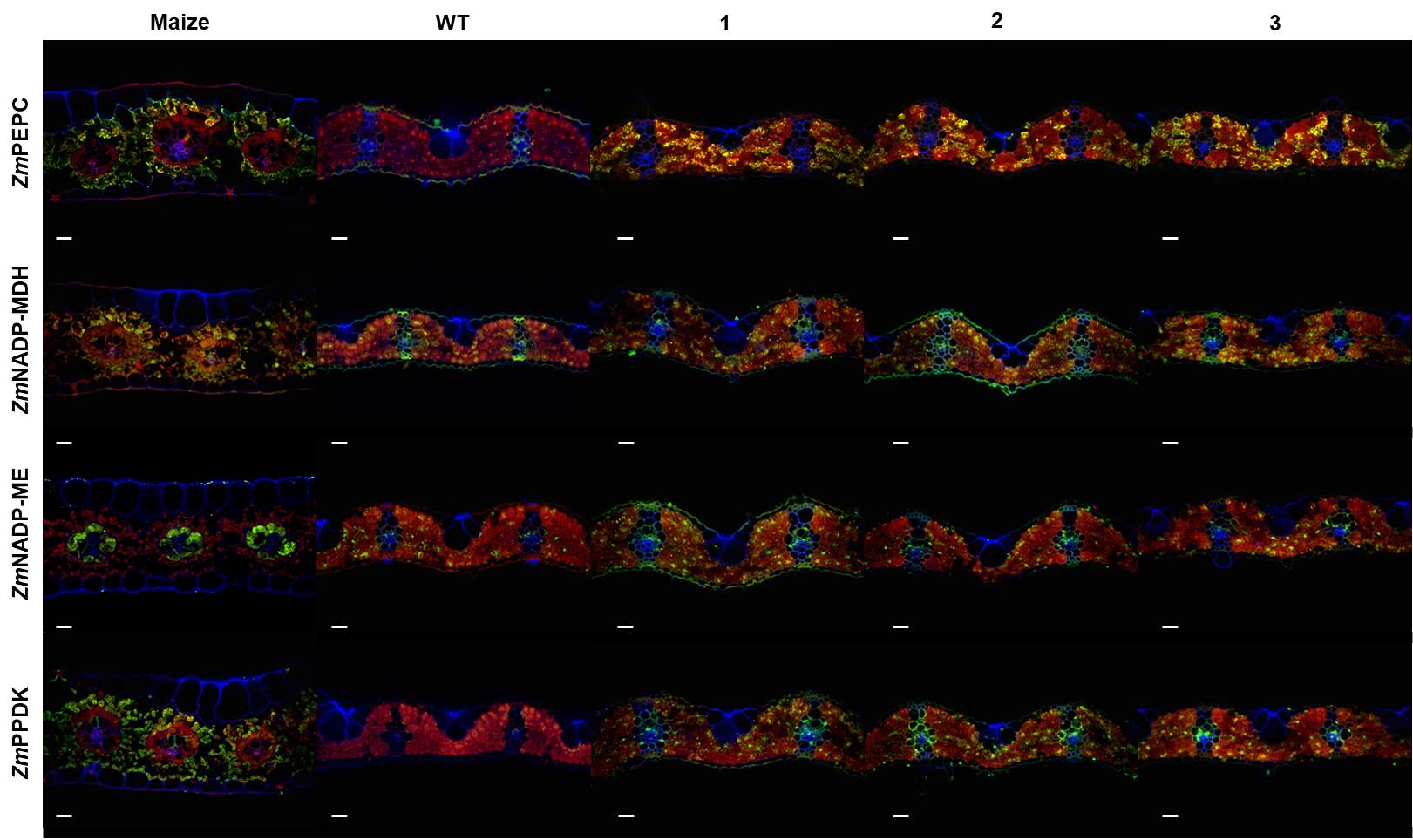

**Supplementary Figure 3.** Representative images of immunolocalization of *Zm*PEPC, *Zm*NADP-MDH, *Zm*NADP-ME and *Zm*PPDK protein. Anti-rabbit polyclonal antisera (*Zm*PEPC 1:500, *Zm*NADP-MDH 1:10, *Zm*NADP-ME 1:100, *Zm*PPDK 1:25) plus Alexa Fluor 488 goat anti-rabbit IgG as secondary antibody (1:200; shown in green color). Red shows autofluorescence of chlorophyll in chloroplasts. Co-staining with calcofluor white visualized cell walls (shown in blue). Scale bar: 20 µm. Images are of the middle portion the seventh fully expanded fifth leaf of maize, wild-type IR64 (WT) and three representative plants for F_2_ generation quadruple crosses. Maize: positive control. Wild-type: negative control.

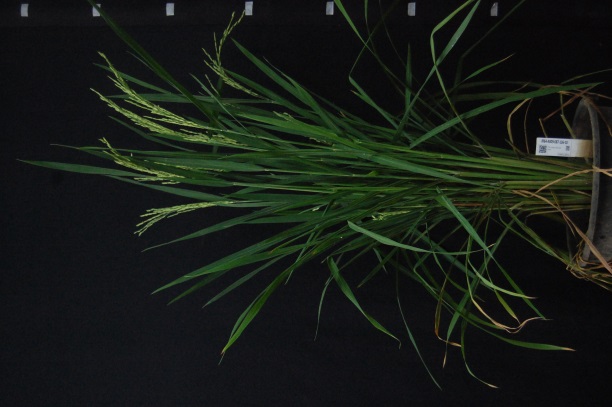

**WT**

**A**

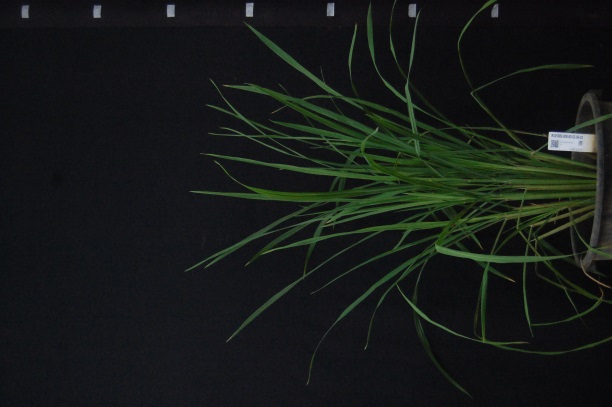

**1**

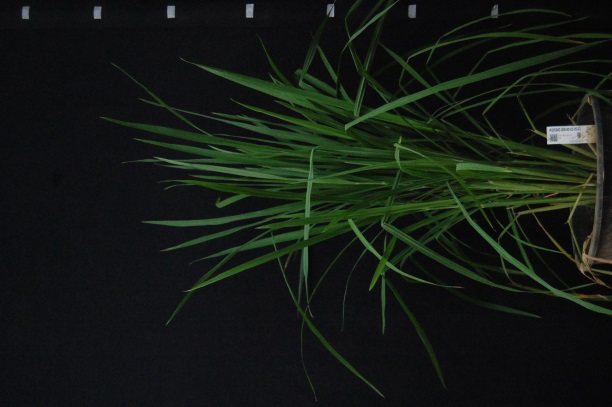

**2**

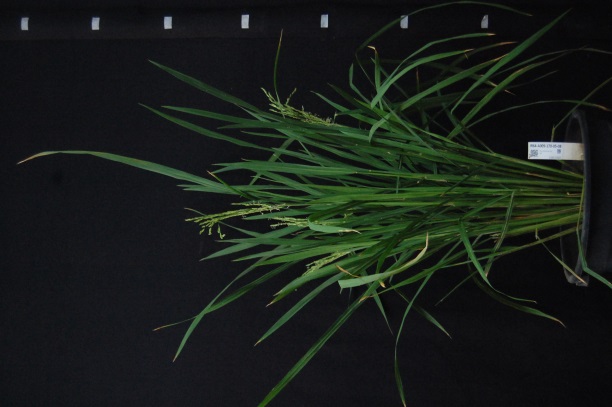

**B**

**WT**

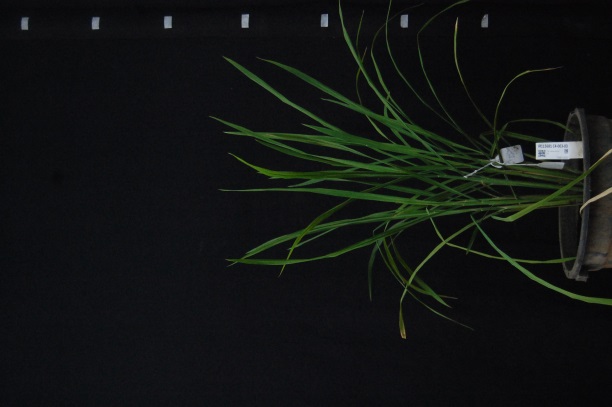

**1**

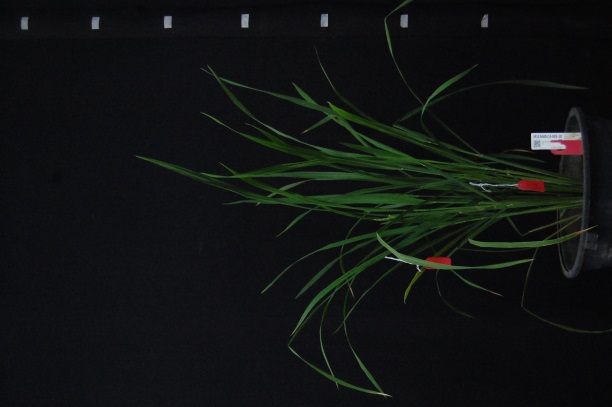

**2**

**Supplementary Figure 4.** Representative pictures of wild-type (WT) and two representative **(A)** quadruple and **(B)** quintuple crosses. Scale bar 10 cm. 90 days post germination (DPG).

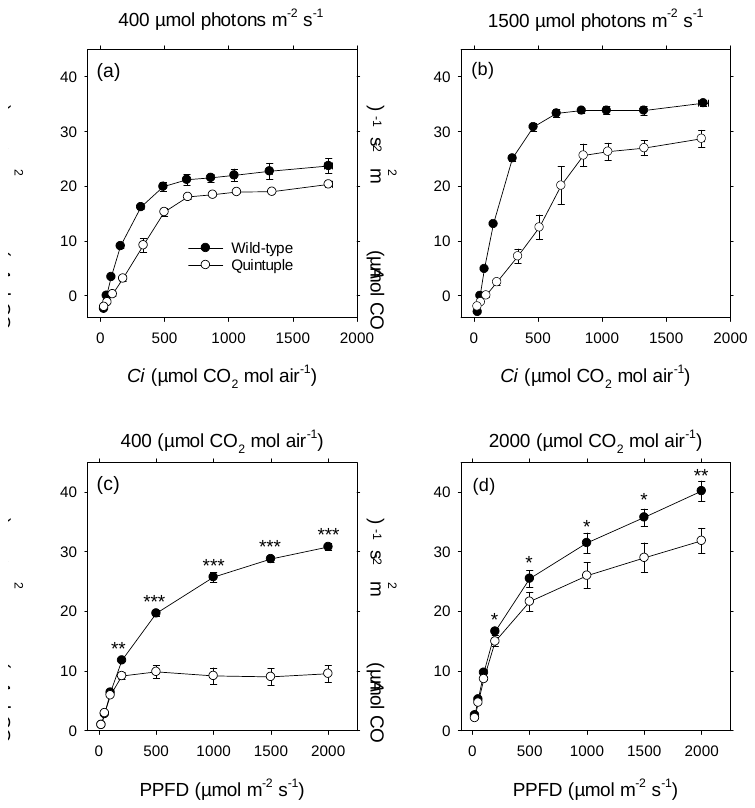

**C**

**A**

**D**

**B**
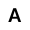

**Supplementary Figure 5. (A, B)** Net CO_2_ assimilation rate (*A*) in response to intercellular *p*CO_2_ (*Ci*,) and (c, d) photosynthetic photon flux density (PPFD). Measurements were made at 400 µmol photons m^-2^ s^-1^ **(A)** and 1500 µmol photons m^-2^ s^-1^ **(B)** under 21 % O_2_. Measurements were made at a *p*CO_2_ (*C_a_*) of 400 µmol CO_2_ mol air^-1^ **(C)** and 2000 µmol CO_2_ mol air^-1^ **(D)**. Values are means ± SE of three individual F_2_ generation quintuple crosses and four wild-type plants. A Student’s t-test was performed for **(C, D)**. Significant differences between WT and quintuple within PARi level are indicated by *P-value<0.05, ** P-value<0.01, *** P-value<0.001.

**Supplementary Table 2.** Comparison of photosynthetic parameters.

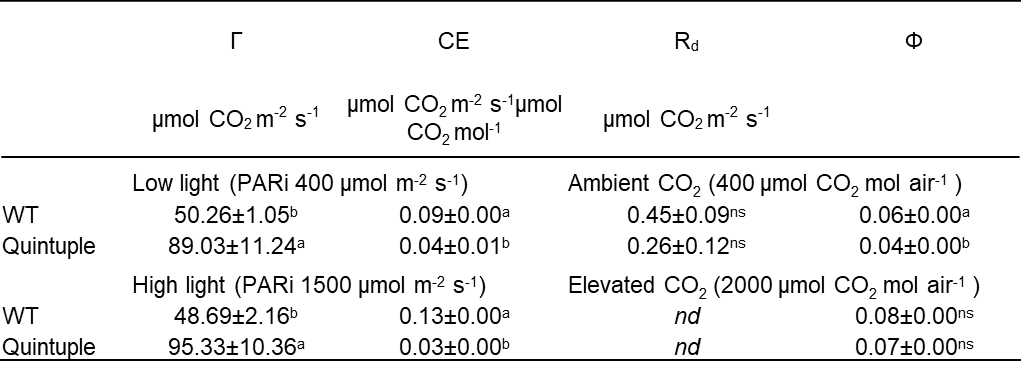

CO_2_ compensation point (*Γ*), carboxylation efficiency (CE), respiration rates (*R*_d_) and quantum yield for CO_2_ assimilation (Φ). Measurements of Γ and CE were made at a PPFD of 400 or 1500 µmol photons m^-2^ s^-1^. Measurement of R_d_ and Φ were made at a *p*CO_2_ (*C_a_*) of 400 or 2000 µmol CO_2_ mol air^-1^ and a leaf temperature of 25°C. Values are means ± SE of three F_2_ generation cross plants and four wild-type (WT) plants. Different letters indicate those values that are statistically different between WT and quintuple within *p*CO_2_ level or PARi level based on a Student’s t-test, P-value<0.05. ns indicates non-significant. *nd*, not determined.

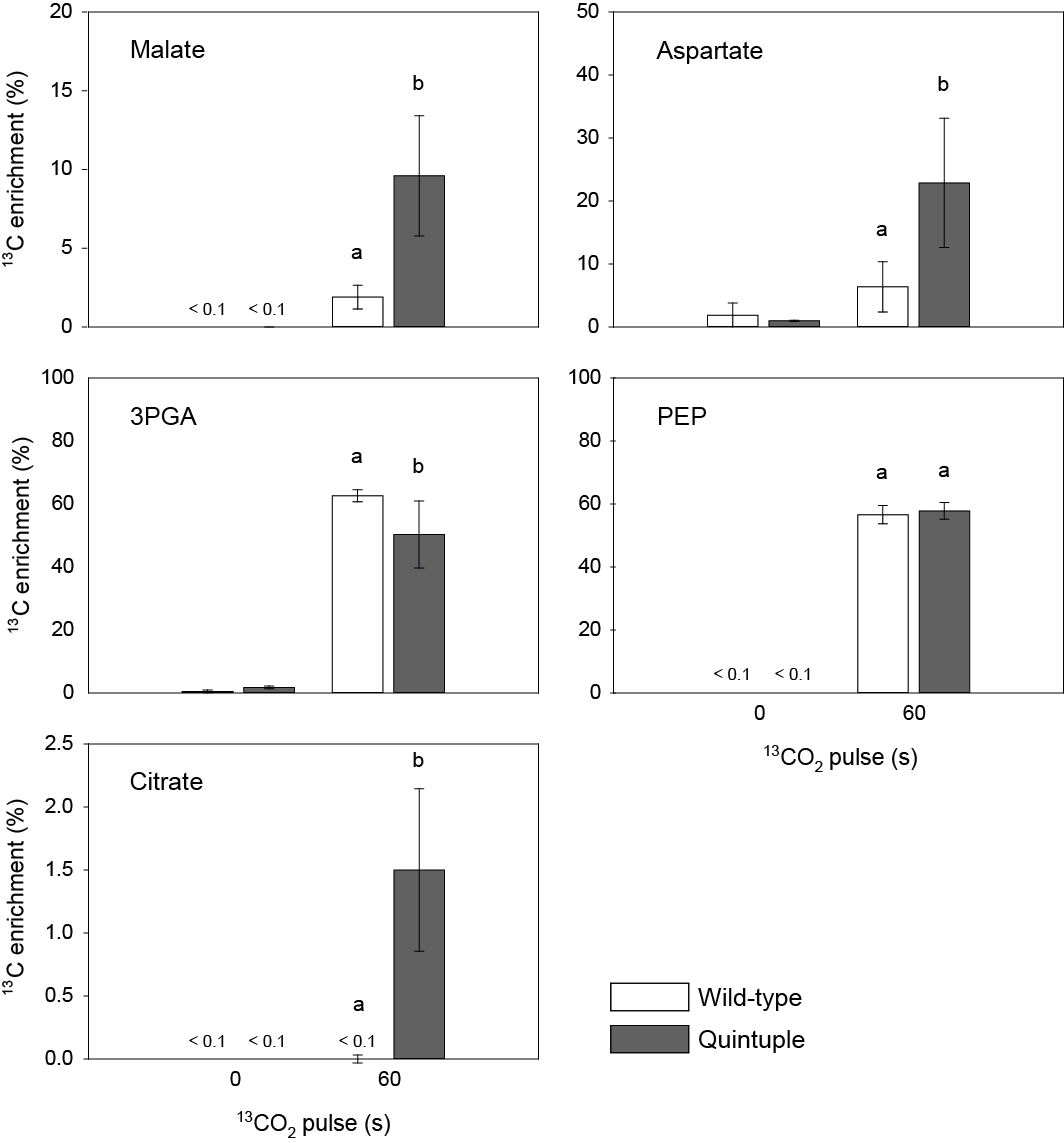

**Supplementary Figure 6.** ^13^CO_2_ pulse-labelling of wild-type and quintuple rice lines. Fully expanded leaves were pulse-labelled with ^13^CO_2_ (300 ppm) for 60 s under steady state photosynthetic conditions. Isotopomers of malate, aspartate, 3PGA, PEP and citrate were measured in extracts from pulse-labelled (60 s) and non-labelled (0 s) leaves by LC-MS/MS, and ^13^C enrichment (%) was calculated after correction for natural abundance. Values are means ± SE of four individual F_2_ generation quintuple crosses and four wild-type plants. The original data are presented in Supplementary Dataset B. Different letters within groups indicate those values that are statistically different based on a one-way ANOVA, P-value<0.05.
